# Patterns of discursive interactions between students and teachers in the biology learning process in High school: the use of social media

**DOI:** 10.1101/2020.04.09.034447

**Authors:** Scarlet Ferreira de Souza, Bárbara Rodrigues Cintra Armellini, Alexandre La Luna, Ana Carolina Ramos Moreno, Martha Cristina Motta Godinho Netto, Rita de Cássia Café Ferreira, Flávio Krzyzanowski Júnior

**Author notes:** Corresponding author: Flávio Krzyzanowski Júnior. Instituto Federal de Educação, Ciência e Tecnologia de São Paulo, São Paulo, SP, Brazil. Vaccine Development Laboratory, Department of Microbiology, Institute of Biomedical Sciences II, Universidade de São Paulo. Conflict of interest disclosure: The authors declare no conflicts of interest.

## Abstract

Increasingly, blending teaching has become a reality in a generation where the digital language is present in virtually every activity. In addition to allowing greater independence and encouraging students to learn at their own pace, blending teaching allows the student to easily access reliable information quickly. Therefore, new studies related to active learning methodology are fundamental. In this study we analyzed 69 interactions between high school students and their teachers in a biology learning activity using a social networking site and the methodology proposed by Mortimer and Scott. The results showed that the prior knowledge of students as well as questions posing challenges and problems to be solved, a very important approach in learning Science Methodology, were barely explored by teachers and mediators (17% and 1%, respectively). Our data demonstrated that the use of digital technology alone does not guarantee interactions that contribute to the learning process in the field of natural sciences. Proposals were also discussed so that these interactions become more diversified and interesting for students, arousing interest in research and promoting the knowledge of scientific methodology.

## Introduction

Internet access has been growing on a daily basis, both in Brazil and throughout the world. From 2015 through 2017, a United Unions research showed an increase in the use of Broadband Internet in the world, where countries such as Brazil, India, and China accounted for approximately 70% of the access [20]. Brazil is the 4th in Internet access and 3rd in terms of time spent accessing the net, according to the website “we are Social” (https://wearesocial.com/). Interestingly, about 62% of the Brazilian population is connected to social networks, which demonstrates its great importance in the national territory. Of all social networking websites, Facebook® stands out with around 2.3 billion users worldwide, with 127 million in Brazil, according to data from the site itself. Therefore, the Internet needs to be considered when thinking about Brazilian education, and specifically, the use of social media, such as Facebook^®^, can be of great value in the development of educational activities.

Regarding this matter, there is a methodology known as Blending Teaching, which mixes traditional classes with the use of virtual environment. In this teaching approach, the main focus ceases to be the teacher, and students become responsible for their learning [1]. This methodology has been studied and debated for a long time, being inserted, in 2003, among the ten main research trends in teaching, revealing its innovative role, inserting the virtual environment in the student learning process [3]. It is important to point out that in the Blending-teaching approach the teacher becomes a mediator of the teaching-learning process, guiding students in the search for answers, encouraging them to “learn to learn”, which was considered by Delors [11] as one of the pillars of education of the 21st century. Nevertheless, most teachers do not use digital technology in their classroom activities [9],which is reflected in the National Curriculum Parameters [7], where digital technologies are seen as merely a tool to assist the individuals in their daily activities. The same can be observed with the National Common Curricular Base (NCCB), launched in 2017 by the Ministry of Education [5]. A research conducted in 2011 showed that about 80% of Brazilian schools had computer labs, but only 22% of teachers used this technology. By 2014, the number of teachers using technology had more than doubled (46% [6]). Despite all this scenario, little is discussed about the use of the Internet in Science Teaching, and even less in Microbiology teaching. Regarding Microbiology, this field of science is acknowledged to have great influence in the flow of life and, therefore, it is important to point out to students how they can benefit from the study of microbiology [8]. News related to microorganisms usually reflects their negative side, such as diseases, which makes students fear these living beings [19]. The current activities and materials proposed in the classroom may not fully collaborate, since the information can be so abstract that it leads to learning difficultly. Therefore, the use of technologies may help, making the approximation of the micro and macroscopic world possible, establishing relations of cause and effect between them [12].

A project entitled “Adopt a Bacteria” [4], which aimed to teach Microbiology in Higher Education, was developed at the Institute of Biomedical Sciences of the University of São Paulo (ICB-USP), in the Microbiology department. Authors used Facebook^®^ as a tool for undergraduates to post articles, videos and news on previously chosen microorganisms [16]. Given the positive results obtained in Higher Education, this approach was applied in the discipline of Biology in High School, expanding the repertoire of microorganisms used (including viruses and fungi), with the name of “Adopt a Microorganism”. The impact of the expansion of the classroom to a virtual environment in teacher-student and student-student interactions as a fundamental base for the teaching and learning process has not yet been analyzed in the literature [21]. It would be possible to assume that the patterns of discursive interaction established in social networking sites would present their own quality and dynamics since space and time are used by teachers and students in a different way from what occurred in the classroom. Thus, the present work seeks to verify the interaction patterns of the activity “Adopt a Microorganism” in a class of 2nd year of High School at the Technical Teaching in Integrated Mechanics of the Federal Institute of São Paulo, in order to analyze if the use of social media as a learning tool guarantees the quality of the interactions, using the analytical approach proposed by Mortimer and Scott [15].

### Study Design

This study was performed at the Federal Institute of Education, Science, and Technology of São Paulo, Campus São Paulo (IFSP) with students from the 2nd year of Technical Education in Mechanics as part of the Biology discipline. The project “Adopt a Microorganism” was carried out during the third quarter of the 2016 school year, with the voluntary participation of 18 students. The study design was performed according to Piantola et al, 2018, as summarized in Figure 1.

**Figure 1.**
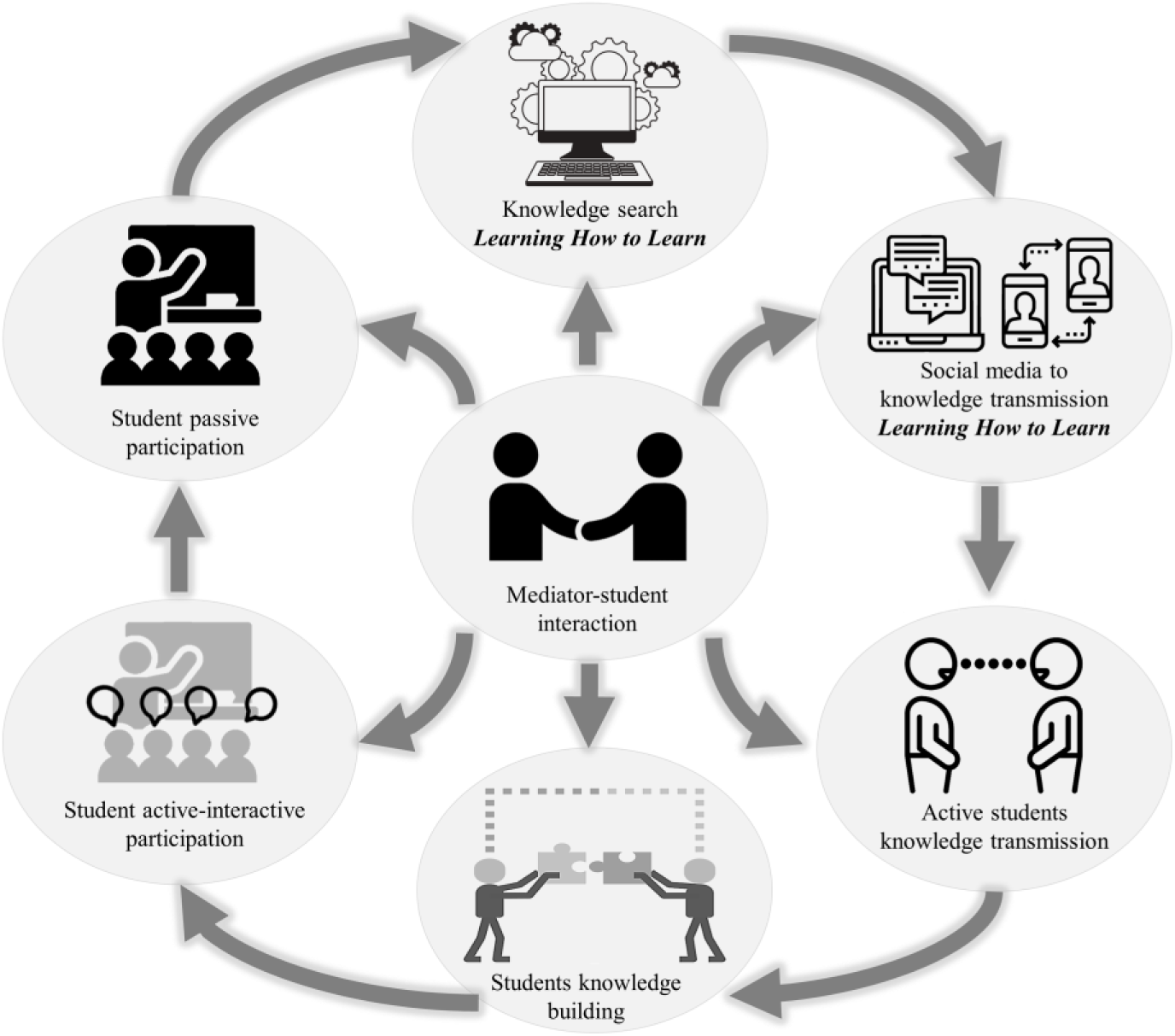
Study design. The educational approach of the “Adopt a Microorganism” is based on four pillars: teacher, mediators, students, and social media

The students of Technical Education in Mechanics were divided into six different groups and each group “adopted” a microorganism (Table 1). Different from those authors who addressed only bacteria, in our study we approached different types of medically important microorganisms. The role of teachers in this system was, apart from conducting the presential classes, also to create challenging questions in order to guide the students through the discussions on Facebook. Every week teachers posted a new challenging question about the adopted microorganism. Each student needed then to answer the question and comment on the posts of colleagues during that week. Students then began to post their own comments about the adopted microorganism. In this scenario, mediators, who were students of graduation in Biology also from the Federal Institute of Education, Science, and Technology of São Paulo had the important and essential role of assisting students in their posts, asking questions and providing the enrichment of the thematic discussions. Consequently, students interacted with both mediators and colleagues, generating a teaching-learning environment where students were the protagonists. Teachers also participated in discussions. Therefore, students began to interact better with their teachers and their classmates, leading to active participation. In this study, only the interactions of one group were analyzed (HIV, with 18 students), and this group presented the biggest number of interactions on Facebook^®^ discussions (69 in total) when compared to the other five groups.

**Table 1.**
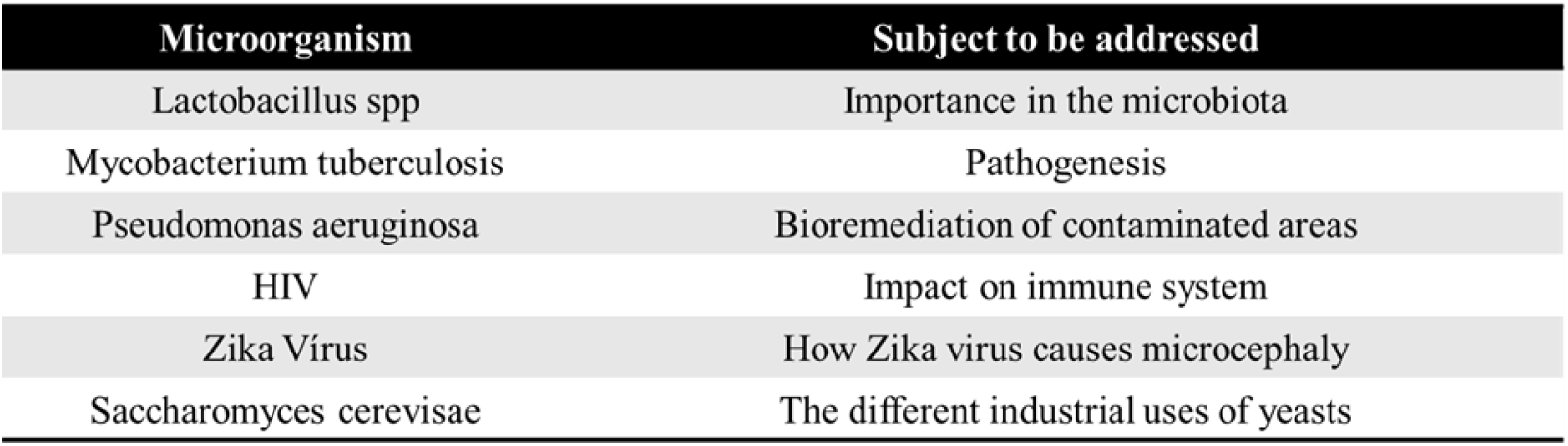
Addressed microorganisms and related topics.

| Microorganism | Subject to be addressed |
| --- | --- |
| Lactobacillus spp | Importance in the microbiota |
| Mycobacterium tuberculosis | Pathogenesis |
| Pseudomonas aeruginosa | Bioremediation of contaminated areas |
| HIV | Impact on immune system |
| Zika Virus | How Zika virus causes microcephaly |
| Saccharomyces cerevisiae | The different industrial uses of yeasts |

### Study development

For the development of this project, some undergraduate students of the Biological Sciences Degree of the IFSP were invited to mediate discussions on Facebook^®^. Before the virtual activities started, the mediators held a workshop with the students to explain how to use reliable sources of scientific information on the internet. Additionally, there were things which were strongly forbidden, such as: offense and criticism of participants, political, sports and religious comments, exposure of personal data and personal matters, inappropriate photos and videos, spams and advertisements. Students, teachers and mediators were instructed to carry out the project activities in their personal electronic devices and private accounts. In addition, during the first week of the project some meetings were held between the students and the teachers in order to clear some doubts about the activities. More importantly, since the use of social media involves the exchanging of personal opinions, which, in turn, might generate unwanted exposure, the entire project was held in a Facebook^®^ secret mode group, in which only the accepted members could visualize its contents. At the end of the project, the students developed publicity material (Posters) regarding their adopted microorganism. These posters were further presented to all communities of the IFSP. Another important role of teachers was to evaluate these posters and the individual participation of the students in the discussions on Facebook^®^ in order to determine a final grade.

### Data analysis

In order to understand the different patterns of discursive interactions in social media among students, mediators and teachers in the biology learning process, we used as reference the tool developed by Mortimer and Scott [14]. Using their approach, we evaluated a triad of analytical structures: Professor/Mediators Intentions, Communicative Approach and Discursive Interactions Patterns, considering these subjects to be the most significant when related to the development of the scientific language and the quality of the interactions performed by the students (Table 2). We also evaluated four types of communicative approach (Table 3) and nine discursive patterns of interactions (Table 4).

**Table 2.**
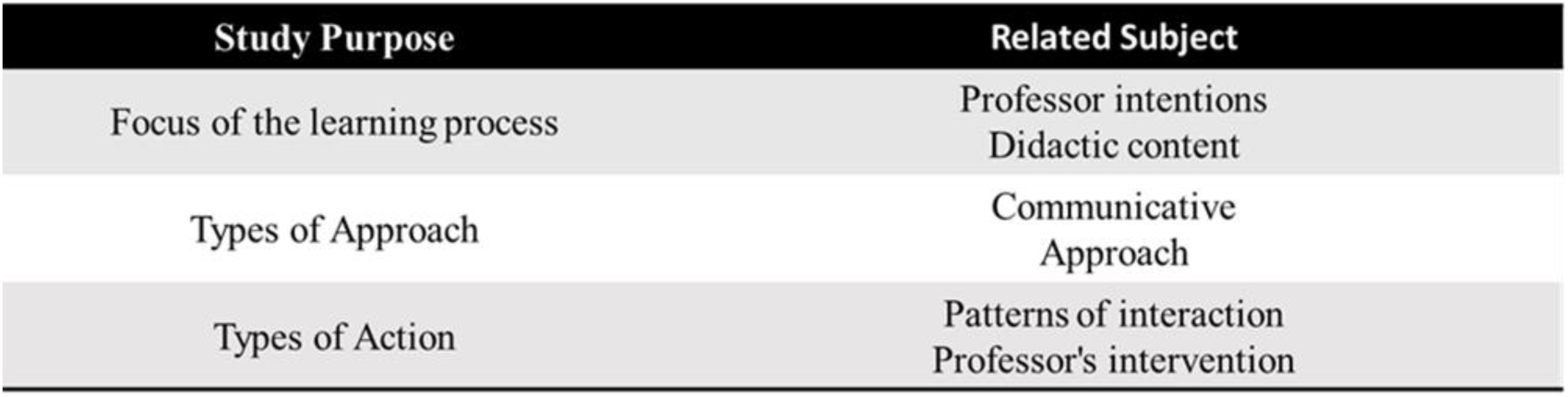
Analytical Structure of the Research Study.

| Study Purpose | Related Subject |
| --- | --- |
| Focus of the learning process | Professor intentions<br>Didactic content |
| Types of Approach | Communicative<br>Approach |
| Types of Action | Patterns of interaction<br>Professor's intervention |

**Table 3.** Types of communicative approach.

| Communicative Approach | Definition |
| --- | --- |
| Interactive / Dialogical | Teacher and students explore and make inquiries about relevant and important issues, from different points of view. |
| Non-interactive / Dialogical | In his speech, teacher reconsiders various points of view, highlighting similarities and differences. |
| Interactive / Authority | Teacher usually leads students through a series of questions and answers, in order to reach a specific point of view. |
| Non-interactive / Authoritative | Teacher presents a specific point of view. |

**Table 4.** Discursive Interaction Patterns: Types of actions in student-mediator interaction.

| Initials | Meaning |
| --- | --- |
| I | Professor/Mediator's initiative |
| SI | Student's initiative |
| SA | Student's answer |
| PF | Professor/Mediator's feedback in order to students elaborate more comments |
| DA | Discursive action that allows continuation of student's comments |
| PE | Professor/Mediator's evaluation |
| LA | Professor/Mediator's Like Assessment |
| RD | Restarting discussion |
| UR | Unrelated response |

With regard to discursive interaction patterns, some adaptations were necessary, since Mortimer and Scott [14] concentrated their attention only in the classical classroom .Thus, in the context of virtual environment, new categories of interaction were required. One of these new categories was “Like Assessment” (LA). Like assessment is present only on Facebook and it is not present in oral interactions. Another adapted category was “Students Initiative” (SI), since the initiative of students was present in this study and had not been predicted in Mortimer and Scott’s model.

In Figure 2 we have an example of the analytical method used to asses the discussions in Facebook^®^ regarding Mortimer and Scott’s approach. For the other categories, the 69 interactions were analyzed following the pattern shown in Figure 3.

**Figure 2.**
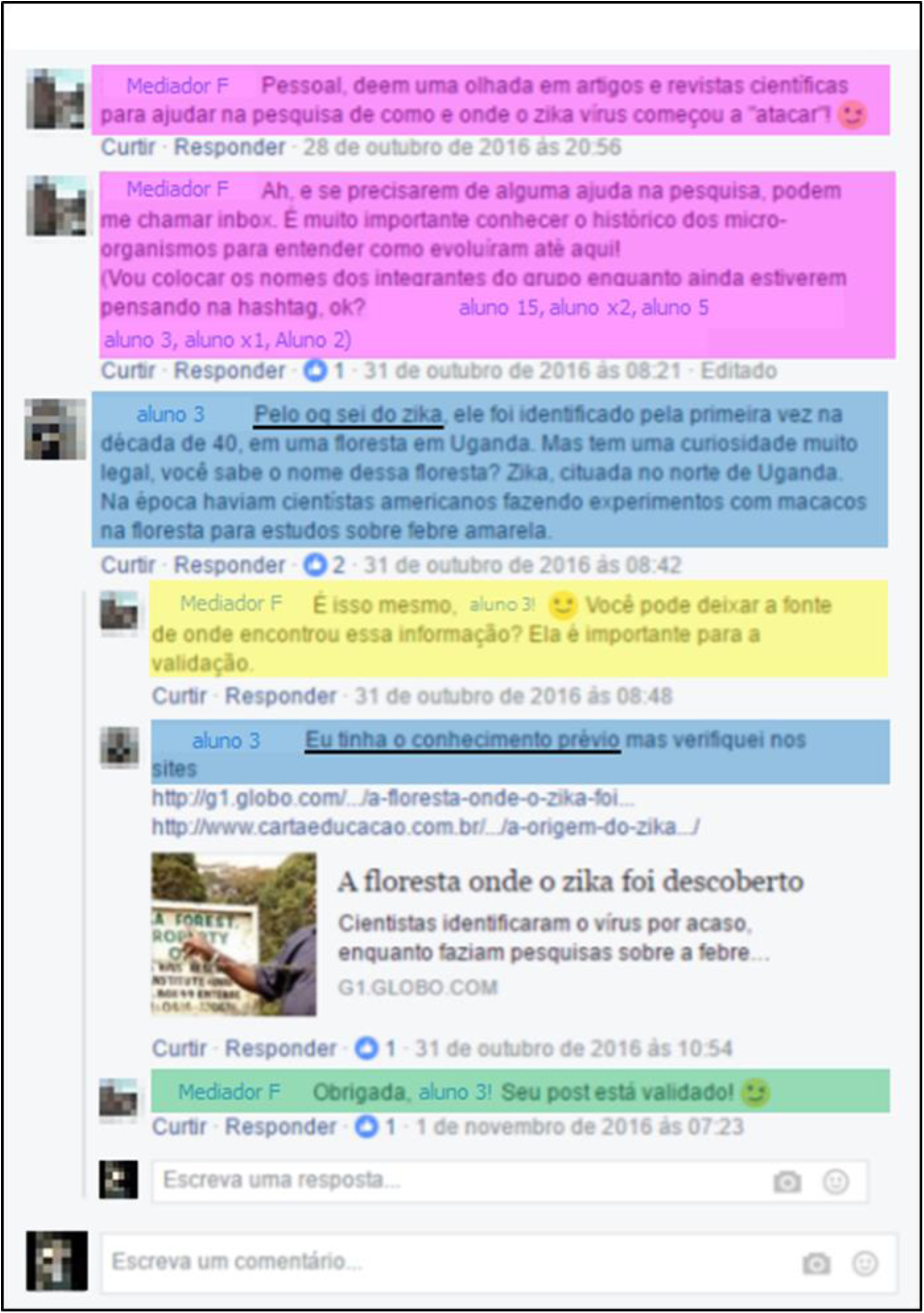
Analytical method used to asses the Discursive Interaction Patterns in Facebook^®^ regarding Mortimer and Scott’s approach. Different colors were used to characterize the 69 different discursive interaction patterns. In this example: Pink represents restarting discussion (RD); Blue, student’s answer (SA); Yellow, discursive action that allows continuations of student’s comments and Green, Professor/Mediator evaluation (PE). Below we present the translation into English:

**Figure 3.**
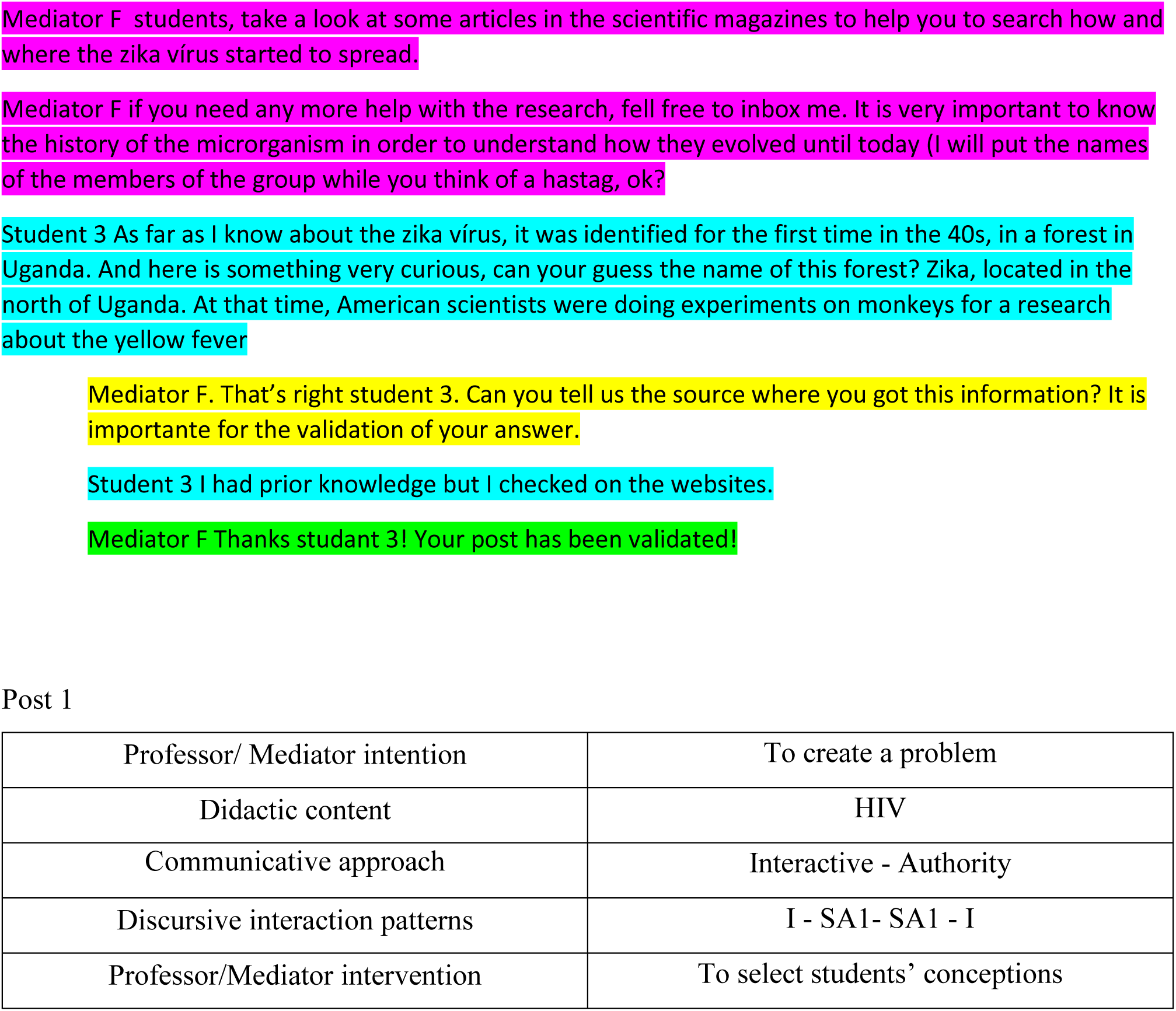
An example of the pattern used to analyse the 69 interactions obtained from Facebook^®^. Only three categories were discussed in this study: Professor/ Mediators Intention, Communicative Approach and Discursive Interaction Patterns.

Mediator F students, take a look at some articles in the scientific magazines to help you to search how and where the zika vírus started to spread.

Mediator F if you need any more help with the research, fell free to inbox me. It is very important to know the history of the microrganism in order to understand how they evolved until today (I will put the names of the members of the group while you think of a hastag, ok?

Student 3 As far as I know about the zika vírus, it was identified for the first time in the 40s, in a forest in Uganda. And here is something very curious, can your guess the name of this forest? Zika, located in the north of Uganda. At that time, American scientists were doing experiments on monkeys for a research about the yellow fever

Mediator F. That’s right student 3. Can you tell us the source where you got this information? It is importante for the validation of your answer.

Student 3 I had prior knowledge but I checked on the websites.

Mediator F Thanks studant 3! Your post has been validated!

## Results

To evaluate the intentions of teachers and mediators during the learning activity on Facebook^®^, the interactions were classified according to the tool developed by Mortimer and Scott [14]. We observed that the items “created problems” and “explore the vision” were little used (1% and 17%, respectively) (Fig. 4). “Keeping the narrative” was the most common intention (81%), evidencing that the intention “creating a problem” was little explored (1%) by professors and mediators. It is noteworthy that this intention (creating a problem) is pivotal in science education.

**Figure 4.**
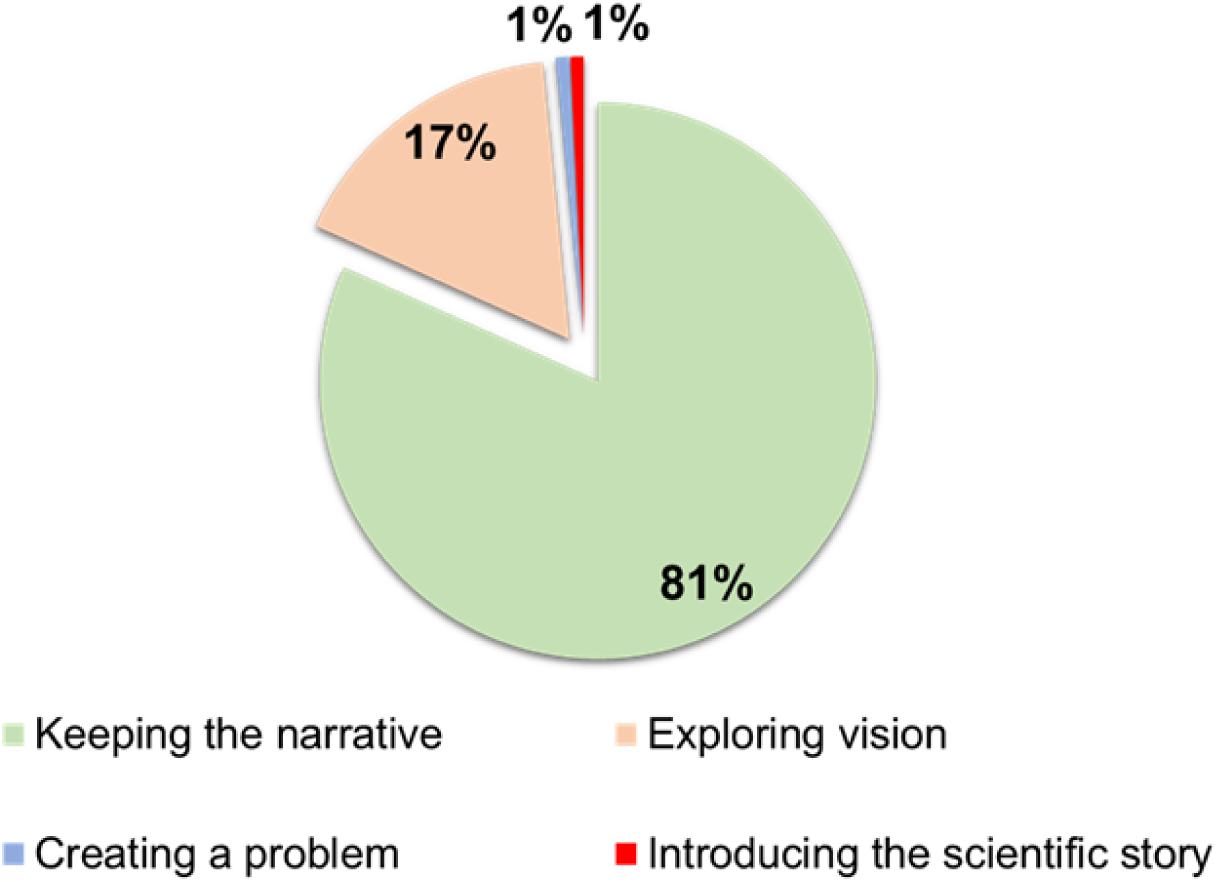
Types of intentions of the discursive action from teachers or mediators.

Regarding the communicative data analysis, most of the interactions were short (I-SA-PE-LA/ I-SA-PE/ SI-SA-PE) (Fig.5), consequently preventing discussions in which the topics that were studied could be deepened, or the prior knowledge of students could be explored. It is important to note that the most appropriate interaction pattern for the student learning process was the least explored by teachers and mediators during the teaching activity (I-RD-SA-PE-DA, 2%).

**Figure 5.**
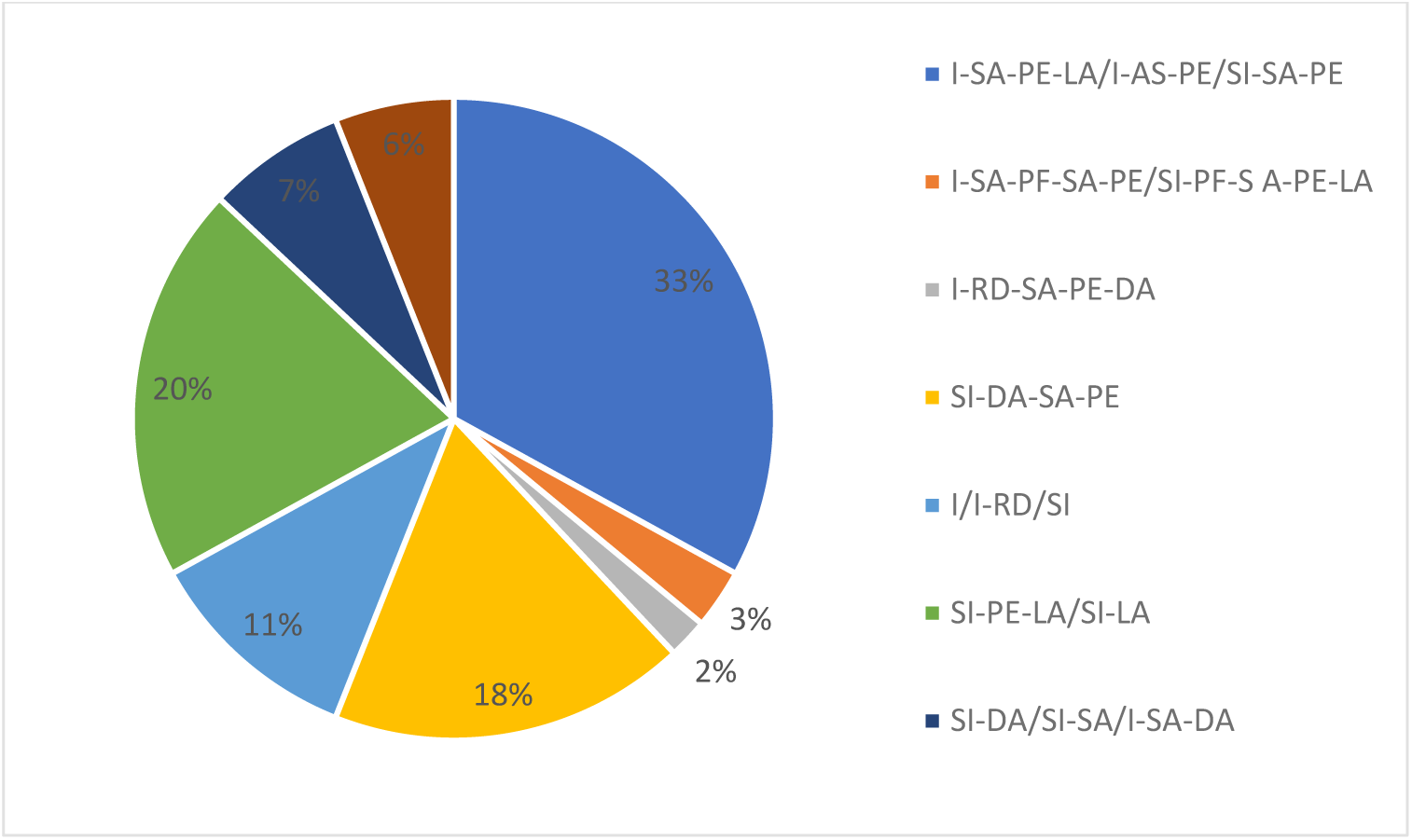
Distribution of the communicative approach found in the 69 interactions among teachers, mediators and students.

## Discussion

In this study we used a discursive interaction analysis tool, created by Mortimer and Scott [15], to evaluate the interactions that occurred in a social networking site during high school biology teaching. In the project “Adopt a Microorganism”, Facebook © was chosen to make teaching viable with a Blending approach [1]. In this approach, actions are taken consciously to allow individualization of teaching. The teacher, seen as the only source of knowledge in traditional teaching, starts to act more as a mediator of the learning process. Activities in which students are protagonists of their own learning are encouraged. One of such activities is encouraging discussions of previously studied items among students during classes.

The social networking site helped promote dialogue and interaction among students and teachers/mediators in a collaborative construction of knowledge. This aspect was also commented by Rabello [17], who also used the Facebook^®^ platform in English language teaching, and analyzed the dynamics of the activity from the socio-historical conceptions of Vygotsky and Bakhtin.

Although the results show that the use of social media allowed students to interact with one other and with the mediators, Mortimer’s tool demonstrated that several adjustments need to be made in the next versions of the project in order to make the interactions more effective. Furthermore, important elements of scientific language such as argumentation, hypothesis gathering and problem-solving should be more elaborated during the discussions. We will, at this point, elucidate points that need to be adjusted.

With regard to the intentions of the professor/mediators (Fig. 4) we noticed that the items “created problems” and “explore the vision” were little used (1% and 17%, respectively). These intentions are essential when we speak of science teaching. Being aware of the prior knowledge of the student is fundamental in teaching practice because it enables the teacher to draw action plans for the acquisition of knowledge up to the level previously intended by the teacher [22].

In Bloom’s scale, problem-solving composes the higher levels that allow for more meaningful learning, preparing the student to solve problems by applying the knowledge gained in practical questions [10].

Considering the types of communicative approaches, the interactive/authority pattern was the most prevalent, being found in 66 of the 69 interactions (data not shown). This pattern reveals the intention of the mediator or teacher to reach a pre-determined target without taking into consideration the different points of view of the students. This obtained pattern could be explained by the fact that students’ previous knowledge was not explored, as shown in figure 2, where the exploring vision item was not very representative (17%).

Mortimer suggests that the approaches should vary, allowing both the mediators and the students to have an active voice in the interactions. Vygotsky [21] states that learning occurs in interactions. Therefore, the more space is given for students to express their opinions, the longer they will have to talk to one other and search for information themselves in reliable sources other than teachers. In so doing, there will be a great contribution in order to reach the variations suggested by Mortimer.

Taking into account the communicative approaches (Fig. 5), we observed that most of the discussions were initiated by students, making the adaptation of pattern I necessary (Initiator of the mediator), including the new term SI (Student Initiation). The student’s initiation was also described in other works that used this tool, such as that of Rezende and Ostermann [18] who carried out research in an online discussion forum, and also in the work of Mortimer et al [14], carried out in the classroom. The initialization of speech by students occurred in fifty-eight (58) dialogic interactions during the project, corresponding to eighty-four percent (84%) of the initiations.

This fact is relevant because it can signal the interest of the student in the subject being addressed as well as in the use of the tool. We can still explore the fact, already observed by other authors, that students who do not manifest themselves in classrooms participate more when meetings occur outside this environment [12]. This finding is very important because it shows that some students begin to interact better in extra-class environments.

Other adjustments, besides the establishment of speech shift SI (student initiation), were necessary for the tool, considering it was created to analyze face-to-face interactions and not asynchronous virtual interactions. One of these adjustments was to institute the acronym LA (like assessment), which was the evaluation given when a professor/mediator accepted what had been published, and the initials RD (restarting discussion) since, unlike a face-to-face lecture, the discussion could restart hours after a post.

Mortimer and Scott [15] indicate that the most common interaction patterns are the triads I-SA-PE (professor/mediator’s initiation, student answers, professor/mediator’s evaluation) that emerge from the alternation of mediator-student speech. As exposed in figure 3, this pattern of classical interaction was evidenced in only fifteen (15) interactions of a total of eighty-four (84) interactions in the sixty-nine (69) postings. However, when I-SA-PE-SI, SI-AS-PE or I-SA(n)-PE are considered to be I-AS-PE derived patterns, the variation of the number of student responses before the mediator evaluation is evident: twenty-eight (28) publications with these interactive standards corresponding, therefore, to thirty-three percent (33%) of the interactions, that is, 33% of the interactions gravitated on the I-AS-PE triad and its variations. These patterns, which are very common in a traditional classroom, show mechanistic participation of the student, and it is then assumed that only through an expositive class, where students should answer questions made by teachers, can they learn.

Rarely did mediators provide feedback for students demanding that they elaborate their answers a little more. The interaction patterns I-SA-PF-SA-PE or SI-PF-SA-PE-SI appeared only twice during the activity, representing only three percent (3%) of interactions of this type. The scarcity of these patterns highlights the fact that mediators were not willing to prolong the discussions, contributing to the impoverishment of the interactions and, consequently, less learning. This would be an important opportunity for mediators to propose the development of argumentation and the resolution of a real problem, using questions that would challenge their prior knowledge.

The second most expressive interaction pattern corresponded to the speech shifts, when the student started a discussion and soon after the mediator evaluated the content of the student’s speech. This interaction was expressed by the acronym SI-PE-LA (students initiative, professor/mediators evaluation, like assessment) or SI-LA, and represented twenty percent (20%) of the total interactions. The student initiation (SI) followed by a professor/mediator evaluation (PE) without the interaction of the other students involved in the project may indicate a low interaction rate in this percentage of postings.

Actions aimed at lowering this pattern are essential in order to increase interactivity. An important suggestion would be for the evaluation to occur after a long discussion. Another suggestion would be to involve other students in ongoing discussions, so that interactions are enriched with the greater participation of students

The data show that only the use of a technological tool, by itself, does not guarantee that important elements in the teaching-learning process, such as the variation of the interactions and their quality, occur naturally [2]. From the point of view of science teaching and its language, the same was observed. This implies that teachers should be trained not only for the use of new technological tools but also for the pedagogically effective use of them.

Having all that was presented in mind, we emphasize that the items that need to be better worked with the professor and mediators in the next versions of the project “Adopt a Microorganism” should be: prolonging discussions with students, so as to find out more about their previous knowledge; a greater variation of interactions, leading to students being able to express their opinions and hypotheses more freely; increasing the use of speech shifts DA (discursive action that allows continuation of student’s answers) and PF (professor/mediator’s feedback), which would allow mediators to present problems in which students would have to apply the acquired knowledge; finally, the extension of the evaluation PE (professor/dediators evaluation) and LA (like assessment) by the professor/mediators. When these goals are achieved, an environment of richer interactions will be provided.

We believe that training mediators to work with the skills and competencies related to the development of argumentation, as well as solving problems, would be a great contribution for the natural enrichment of discussions, as well as developing the scientific language in students.

## Notes

### Competing Interest Statement

The authors have declared no competing interest.

## References

1. Bacich L, Neto AT et al (2015) Ensino híbrido: personalização e tecnologia na educação. Penso Editora. Porto Alegre.

2. Barroqueiro CH, Bonici R et al (2000) O uso das tecnologias de informação e comunicação no ensino de ciências e matemática: uma bênção ou um problema? Encontro nacional de pesquisa em educação em ciências. Florianópolis.

3. Bonk J, Graham CR (2012) The handbook of blended learning: Global perspectives, local designs. John Wiley & Sons

4. Botte DAC, Souza RD et al (2014) Microbiologia no ensino superior: “adote uma bactéria!” (e o facebook©)!. Microbiologia In Foco. Rev Microbiol São Paulo, v. 23, p.5–9, Ano 5

5. Brasil (2017) Base Nacional Comum Curricular. Brasília: Ministério da Educação.

6. Brasil (2014). INEP. Instituto Nacional de Estudos e Pesquisas Educacionais Anísio Teixeira. Brasília: Ministério da Educação.

7. Brasil (1999) Parâmetros Curriculares Nacionais. ensino médio. Brasília: Ministério da Educação.

8. Cassanti AC, Araújo EE et al (2008) Microbiologia democrática: estratégias de ensino-aprendizagem e formação de professores. Revista Conhecer, v. 9, n. 1, p. 84–93

9. Corrêa HT, Dias D (2016) Multiletramentos e usos das tecnologias digitais da informação e comunicação com alunos de cursos técnicos. Trab Linguist Apl vol.55, n.2, pp. 241–262

10. Crowe AD, Wenderot NP (2008) Biology in Bloom: Implementing Bloom’s taxonomy to enhance student learning in Biology. CBE Life Sci Educ Vol (7): 368–381

11. Dellors J et al (1999) Educação: um tesouro a descobrir: relatório para a UNESCO da Comissão Internacional sobre Educação para o Século XXI. Educação: um tesouro a descobrir: relatório para a Unesco da Comissão Internacional sobre Educação para o Século XXI

12. Grosseck G, Bran R et al (2011) Dear teacher, what should I write on my wall? A case study on academic uses of Facebook. Procedia Soc Behav Sci 15: 1425–1430

13. Kimura AH, Oliveira GS et al (2013) Microbiologia para o ensino médio e técnico: contribuição da extensão ao ensino e aplicação da ciência. Revista Conexão UEPG, v. 9, n. 2, p. 254–267

14. Mortimer et al. (2007) Uma metodologia para caracterizar os gêneros de discurso como tipos de estratégias enunciativas nas aulas de ciências. In: NARDI, R. (Org.). A pesquisa em ensino de ciências no Brasil: alguns recortes. São Paulo: Escrituras. p. 53–94.

15. Mortimer E F, Scott P H (2002) Atividade discursiva nas salas de aula de ciências: uma ferramenta sociocultural para analisar e planejar o ensino. Investigações em Ensino de Ciências, Porto Alegre, v. 7, n. 3, p. 283–306.

16. Piantola M A F et al. (2018) Adopt a Bacterium – an active and collaborative learning experience in microbiology based on social media. Braz J Microbiol vol.49, n. 4, pp. 942–948.

17. Rabello C R L (2015) Interação e aprendizagem em Sites de Redes Sociais: uma análise a partir das concepções sócio-históricas de Vygotsky e Bakhtin. Rbla, Rio de Janeiro, v. 15, n. 3, p.735-760.

18. Rezende F, Ostermann F (2006) Interações discursivas on-line sobre Epistemologia entre professores de Física: uma análise pautada em princípios do referencial sociocultural. Revista Electrónica de Enseñanza de las Ciencias, v. 5, n. 3, p. 505–521.

19. Sockett Liz (2001) Microbiology-A lifetime’s education. Microbiology Today, v. 28, p. 51–51

20. United Nations (Org.). Information Economy Report (2017): Digitalization, Trade and Development. Nova Iorque e Genebra: United Nations, p. 111

21. Vygotsky L S A (1991) Formação social da mente: o desenvolvimento dos processos psicológicos superiores. 4^a^ ed. São Paulo: Martins Fontes.

22. Zabala A (1998) A prática educativa: como ensinar. São Paulo. Ed. Artmed.

